# Deep convolutional neural networks for accurate somatic mutation detection

**DOI:** 10.1101/393801

**Authors:** Sayed Mohammad Ebrahim Sahraeian, Ruolin Liu, Bayo Lau, Marghoob Mohiyuddin, Hugo Y.K. Lam

## Abstract

We present NeuSomatic, the first convolutional neural network approach for somatic mutation detection, which significantly outperforms previous methods on different sequencing platforms, sequencing strategies, and tumor purities. NeuSomatic summarizes sequence alignments into small matrices and incorporates more than a hundred features to capture mutation signals effectively. It can be used universally as a stand-alone somatic mutation detection method or with an ensemble of existing methods to achieve the highest accuracy.

## Main Text

Somatic mutations are critical signatures in cancer genesis, progression, and treatment. Accurate detection of somatic mutations is challenging due to tumor-normal cross contamination, tumor heterogeneity, sequencing artifacts, and coverage. In general, effectively filtering false-positive calls, which are introduced by the aforementioned issues, and precisely keeping hard-to-catch true-positive calls, which may occur with low allele-frequency (AF) or occur in low-complexity regions, are crucial for an accurate somatic mutation detection algorithm.

To date, a range of tools have been developed to address the somatic mutation detection problem including MuTect2 [1], MuSE [2], VarDict [3], VarScan2 [4], Strelka2 [5], and SomaticSniper [6]. These tools employ different statistical and algorithmic approaches, which perform well in certain cancer or sample types for which they were designed; however, they are limited in generalization to a broader range of sample types and sequencing technologies and thus may exhibit suboptimal accuracy in such scenarios [7–9]. In our earlier work, SomaticSeq [10], we used an ensemble approach to maximize the sensitivity by integrating algorithmically orthogonal methods. It also used machine learning to integrate almost a hundred features to keep the precision high, leading to an accuracy improvement over all individual methods. Nevertheless, the machine-learning backbone used in SomaticSeq relies on a set of extracted features for the mutations’ locations. As a result, it cannot fully capture the raw information in the genomic contexts of the somatic mutations to further distinguish true somatic mutations from background errors, limiting its performance in challenging situations, such as low complexity regions and low tumor purity.

Here we address the limitation of generalizability and complexity of statistical modeling of tumor sequencing data by leveraging deep Convolutional Neural Networks (CNNs). CNNs have recently shown significant performance in classification problems from different domains including germline variant calling [11–13] and skin cancer classification [14]. Even so, applying CNNs to the challenging problem of somatic mutation detection has not been explored. The only previous deep learning based attempt [15] was to apply a six-layer fully-connected neural network to a set of manually extracted features. This approach lacks the power provided by the CNN architecture, which is to learn feature representations directly from the raw data using patterns seen in local regions. Besides, due to the complexity of fully-connected networks, it has less generalizability and scalability as seen in CNNs.

We introduce NeuSomatic, the first CNN-based approach for somatic mutation detection that can effectively leverage signals derived from sequence alignment, as well as from other methods to accurately identify somatic mutations. Unlike other deep learning based methods that are focused on germline variants, NeuSomatic is addressing a bigger unmet need in terms of accuracy due to the complexity of tumor samples. It can effectively capture important mutation signals directly from the raw data and consistently achieve high accuracy for different sequencing technologies, sample purities, and sequencing strategies such as whole-genome vs. target enrichment.

The inputs to NeuSomatic’s network are candidate somatic mutations identified by scanning the sequence alignments for the tumor sample as well as the matched normal sample (Figure 1; Supplementary Figures 1 and 2). Somatic mutations reported by other methods can also be included in this list of candidates. For each candidate locus, we construct a 3-dimensional feature matrix *M* (of size *k ×* 5 *×* 32), consisting of *k* channels each of size 5 *×* 32, to capture signals from the region centered around that locus. Each channel of the matrix *M* has 5 rows representing the 4 nucleotide bases as well as the gapped base (‘-’), and 32 columns representing the alignment columns around the candidate location.

**Figure 1:**
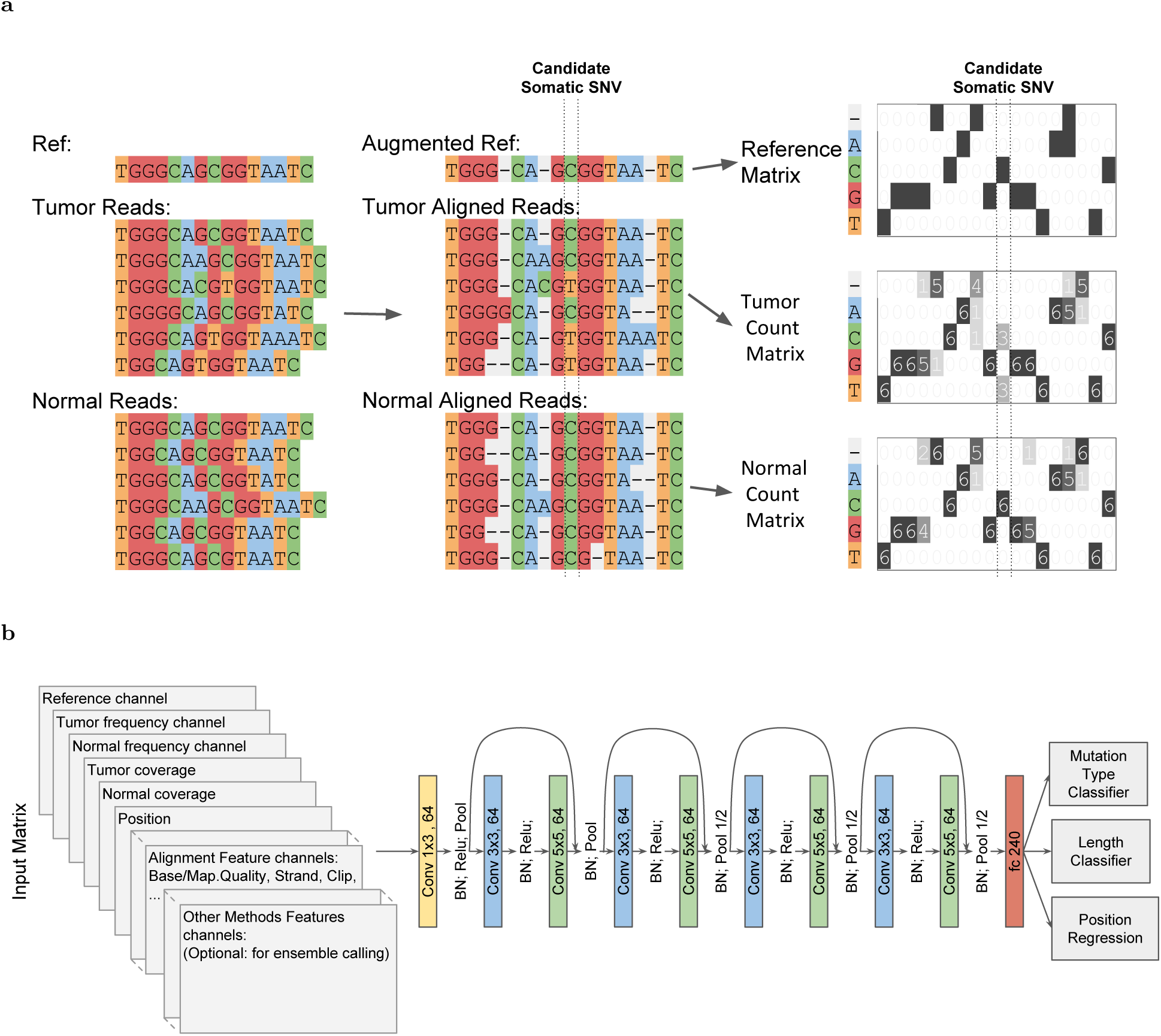
(a) Toy example of input matrix preparation for a given candidate somatic SNV. Sequence alignment information in a window of 7 bases around the candidate somatic mutation is extracted. The reference sequence is then augmented by adding gaps to account for insertions in the reads. The augmented alignment is then summarized into the reference matrix, the tumor count matrix, and the normal count matrix. The count matrices record the number of A/C/G/T and gap (‘-’) characters in each column of the alignment, while the reference matrix records the reference bases in each column. The count matrices are then normalized by coverage to reflect base frequencies in each column. Separate channels are reserved to record the tumor and normal coverages. (b) The input 3-dimensional matrix and the proposed NeuSomatic network architecture. The input matrix consists of reference channel, tumor and normal frequency channels, coverage and position channels, followed by several channels summarizing the alignment features. When used in ensemble mode, NeuSomatic also includes additional channels for other individual methods features. NeuSomatic network architecture consists of 9 convolutional layers structured in four blocks with shortcut identity connections. We use two softmax classifiers and one regressor on the final layer to predict the mutation type, size, and position.

**Figure 2:**
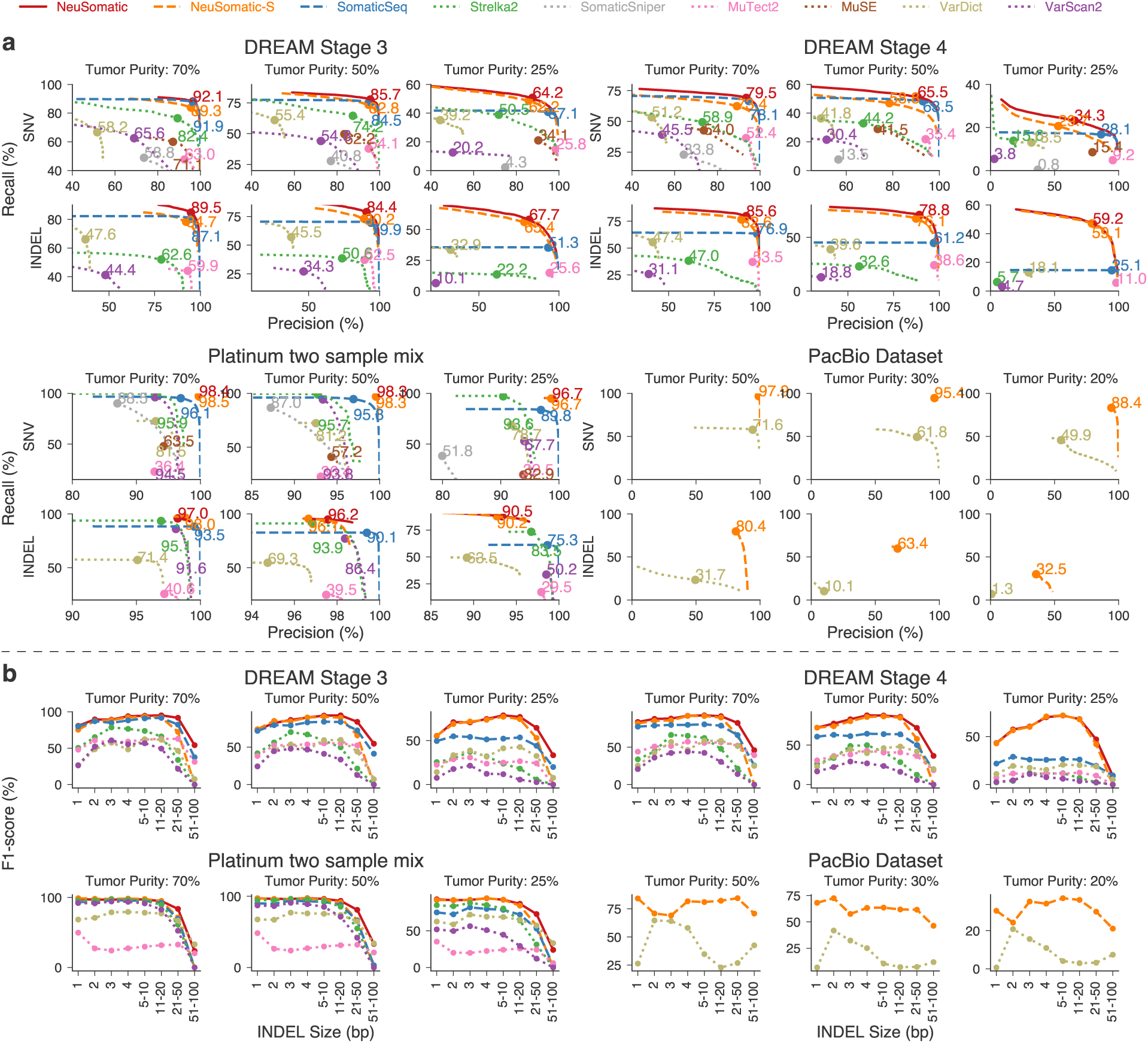
(a) Performance analysis of DREAM Stage 3, DREAM Stage 4, Platinum two sample mixture, and PacBio datasets. For the first three datasets, three tumor purity scenarios (70%, 50% and 25%) are used while normal sample has 95% purity. For the PacBio dataset, three tumor purities (50%, 30%, and 20%) are used while the normal sample has 95% purity. The confidence or quality scores are used to derive the precision-recall curves. The highest F1-score achieved by each algorithm is printed on the curve and marked with a solid circle. (b) Performance analysis of INDEL accuracy (F1-score) for different INDEL sizes.

The first 3 channels, respectively, are the reference, tumor-frequency, and normal-frequency channels that summarize the reference bases around the candidate locus, as well as the frequency of different bases in that region. We augment the reference sequence around the candidate locus with gaps corresponding to the insertions in the read alignments (Figure 1a; Supplementary Figures 1 and 2) in order to capture the insertions. Thus, each column of tumor and normal frequency matrices represents the frequency of A/C/G/T/gap bases in the corresponding multiple sequence alignment (MSA) column of the tumor and normal samples, respectively. The remaining channels summarize other features such as coverage, base quality, mapping quality, strand-bias, and clipping information for reads supporting different bases. If NeuSomatic is used in the ensemble mode, we also use additional channels for features reported by the individual somatic mutation detection methods. With this concise, yet comprehensive structured representation, NeuSomatic can use the necessary information in tumor, normal, and reference to differentiate difficult to catch somatic mutations with low AF from germline variants, as well as sequencing errors. This design also enables the use of convolutional filters in the CNN to capture contextual patterns in the sub-blocks of the matrix.

To compare to other CNN approaches used in genomics problems, DeepVariant [11] uses read pileup as the input for germline variant calling. In contrast, we use base frequency summary for each column as the input to our network. This simplifies the CNN structure, allowing a substantially more efficient implementation. Another germline variant calling method, Clairvoyante [12], uses three channels to summarize base counts for allele counts, deletions, and insertions at the center of the window. In contrast, we summarize all these events in a single base frequency matrix using the reference augmentation approach described earlier, which can clearly represent all the insertion and deletion (INDEL) events across the window.

NeuSomatic employs a novel CNN structure that predicts the type and length of a candidate somatic mutation given the feature matrix *M* (Figure 1b). The proposed CNN consists of 9 convolutional layers structured in four blocks with shortcut identity connections inspired by ResNet [16] but with a different formation to adapt to the proposed input structure. We use two softmax classifiers and one regressor on the final layer. The first classifier identifies whether the candidate is a non-somatic call, SNV, insertion, or deletion. The second classifier predicts the length of the somatic mutation with four classes (0 indicating non-somatic, or lengths from 1, 2 or greater than 2), and the regressor predicts the location of the somatic mutation. Using the output of these classifiers we identify the set of somatic mutations. If the lengths of INDELs are predicted to be larger than 2, we perform a simple post processing step on reads overlapping that position to resolve the INDEL sequence from the read alignment CIGAR string. This has been shown to perform well for data generated by Illumina sequencers. For higher error rate sequencing data, more complex local realignment post-processing is conducted to resolve the INDEL sequence.

Since NeuSomatic can be used in stand-alone and ensemble modes, we use NeuSomatic-S to denote the stand-alone mode, while reserving NeuSomatic to denote the ensemble mode. We compared NeuSomatic and NeuSomatic-S against the state-of-the-art somatic mutation detection methods including MuTect2 [1], MuSE [2], SomaticSniper [6], Strelka2 [5], VarDict [3], and VarScan2 [4], and against the ensemble approach, SomaticSeq [10]. We compared and contrasted performance using multiple synthetic and real datasets. We report below, the synthetic datasets in increasing order of somatic mutation detection difficulty considering the AF of somatic mutation in the datasets.

For the first synthetic dataset, as in previous studies [5, 10] we mixed two normal Platinum Genomes [17] samples, NA12877 and NA12878, at 70:30, 50:50, and 25:75 tumor purity ratios to create three tumor contamination profiles, and at 5:95 ratio to create a contaminated normal sample. We used the germline variants in NA12878 which were reference calls in NA12877 as truth set for the evaluation. Both NeuSomaticS and NeuSomatic significantly outperformed all other methods (Figure 2a and Supplementary Table 1). NeuSomatic’s performance improvement over other approaches increased with lower, more challenging tumor purities (25:75 mixture). In summary, NeuSomatic yielded up to 98.4% and 96.9% F1-scores for SNVs and INDELs respectively overall and an improvement of up to 6.7% over the best method in the lowest sample purity for this dataset.

For the second synthetic dataset, we used the ICGC-TCGA DREAM Challenge Stage 3 and Stage 4 datasets [18], which were constructed by computationally spiking mutations into a healthy genome of a paired normal sample with different AFs (See Methods). We mixed the tumor and normal samples to create three tumor mixtures and a contaminated normal sample with ratios mentioned above. NeuSomaticS outperformed all other stand-alone methods for both Stage 3 and Stage 4 datasets by over 10% for SNVs and 30% for INDELs on average (Figure 2a; Supplementary Tables 2 and 3). This performance improvement increased with decreasing tumor purity (the 25:75 mixture). We further observed that NeuSomatic (the ensemble mode) clearly outperformed both SomaticSeq and NeuSomatic-S, even though NeuSomatic-S still outperformed SomaticSeq in more challenging scenarios, such as SNVs in the 25:75 mixture and INDELs in the 25:75 and 50:50 mixtures. In summary, NeuSomatic yielded up to 92.1% and 89.4% F1-scores for SNVs and INDELs, respectively, overall and an improvement of up to 34.1% over the best method in the lowest sample purity.

For the third synthetic dataset, as in previous studies [1, 10], we constructed a tumor sample by spiking reads from NA12878 into NA12877 in variant positions of NA12878 with spike in frequencies sampled from a binomial distribution with means [0.05, 0.1, 0.2, 0.3] and used an independent set of NA12877 reads as pure normal. Note that, unlike earlier strategy which mixed samples in fixed proportions yielding somatic mutations at fixed AFs, this mixing approach generated them at varying AFs ranging from 0.025 to 0.3. NeuSomatic yielded 80.9% and 66.7% F1-scores for SNVs and INDELs, respectively, over all AFs and an improvement of up to 4% over the best method (Supplementary Figure 3; Supplementary Table 4). For low AF somatic mutations, the performance improvement was even higher (11% improvement for AF=0.025 and 8% improvement for AF=0.05) (Supplementary Figure 3b).

It is worth noting that NeuSomatic consistently outperformed other methods for various INDEL sizes in different datasets (Figure 2b; Supplementary Figure 3c; Supplementary Figure 4). For large (> 50 bases) INDELs, since most of the short reads with somatic INDELs are soft-clipped, the INDEL information is lost in the pileup count matrices. For such cases, NeuSomatic benefited from other methods’ predictions, since some of the methods like VarDict and MuTect2 used local assembly for their predictions.

To assess the performance of NeuSomatic on different target enrichments, we used a whole-exome and a targeted panel dataset from the Ashkenazi Jewish trio [19] (Supplementary Figures 5 and 6; Supplementary Tables 5 and 6). We trained NeuSomatic and SomaticSeq on the whole-exome dataset and applied the trained model on both the whole exome and the panel. For whole exome, NeuSomatic achieved up to 98.1% and 90.9% F1-scores for SNVs and INDELs, respectively. For the targeted panel, NeuSomatic consistently outperformed other methods with 98.8% F1-score for SNVs. Applying the model trained on whole-genome Platinum-mixture data on both target enrichment sets yielded similar performance which confirmed the robustness of NeuSomatic (Supplementary Figures 7 and 8).

We assessed the robustness of NeuSomatic’s training for specific purity by training and testing on different purities for the DREAM Challenge Stage 3 datasets. Supplementary Figure 9 shows that performance degrades only marginally even when we trained and tested on very different tumor purities. We also observed that training using data aggregated from multiple tumor purities was as good as training on individual tumor purities (Supplementary Figure 9). This suggests that a training set incorporating multiple tumor purities is sufficient to get a model which is robust to tumor purity variation.

We further evaluated NeuSomatic’s performance on reads with high error rates, particularly those from the long read sequencing platforms. We used tumor-normal pair samples simulated with 20%, 30%, and 50% AF somatic mutations based on the raw PacBio reads (Figure 2, Supplementary Table 7). NeuSomatic identified somatic SNVs and INDELs with F1-scores of up to 97.9% and 80.4% respectively, which outperformed VarDict [3] by up to 38.5% for SNVs and up to 53.3% for INDELs. This analysis confirms the capability of NeuSomatic in detecting somatic mutations even when the sequence reads have high error rate as in PacBio long raw reads.

For all datasets discussed, we also assessed the performance of INDEL calling by different somatic mutation detection methods using the more relaxed criterion of simply predicting the positions of the somatic INDELs correctly (and ignoring the exact INDEL sequence). Again, we observed similar superiority of NeuSomatic over other schemes indicating that the main improvements are contributed by the proposed CNN structure and not the post-processing INDEL resolution steps (Supplementary Figures 10 and 11).

In the absence of a high-quality, comprehensive ground truth dataset for somatic mutations [20], like the Genome-in-a-bottle gold set for germline variants, we would not be able to calculate F1 accuracy outside of synthetic data. Fortunately, there are existing datasets with validated somatic mutations we could take to estimate the accuracy performance of NeuSomatic in real data (See Methods). We used COLO-829 [21, 22], an immortal metastatic malignant melanoma cell line-derived whole-genome dataset with 454 validated somatic SNVs. To evaluate NeuSomatic on this real WGS sample, we used models trained on the DREAM Challenge Stage 3 with 70% purity. As shown in Supplementary Table 8, NeuSomatic achieved the highest pseudo F1-score of 99.6% for the COLO-829 malignant melanoma sample. We also evaluated NeuSomatic on a TCGA [23, 24] whole exome sequencing (WES) sample of colorectal adenocarcinoma (TCGA-AZ-6601), achieving the highest pseudo F1-score of over 99.6%(Supplementary Table 9).

NeuSomatic is the first deep learning based framework for somatic mutation detection, which is high-performing and universal. While using the same CNN architecture, it achieves the best accuracy for varying tumor purities across multiple datasets ranging from synthetic to real, across multiple sequencing strategies ranging from whole genome to targeted as well as across multiple sequencing technologies ranging from short reads to high-error long reads. Specifically for low tumor purities and low allelic frequencies, NeuSomatic significantly outperforms other state-of-the-art somatic mutation detection methods, thus demonstrating its capability in addressing the hard problem. We believe NeuSomatic advances the state of the art significantly by providing a very broadly applicable approach for somatic mutation detection.

## Online Methods

### ICGC-TCGA DREAM Challenge data

Stage 3 data consists of a normal sample and a tumor sample constructed by computationally spiking 7,903 SNVs and 7,604 INDELs mutations into a healthy genome of the same normal sample with three different AFs of 50%, 33%, and 20% to create synthetic but realistic tumor–normal pairs. Stage 4 data has similar formation, but with 16,268 SNVs and 14,194 INDELs in two subclones of 30% and 15% AF. We then constructed an impure normal by mixing 95% normal and 5% tumor reads. We also constructed 3 tumor mixtures by mixing tumor and normal respectively at 70:30, 50:50, and 25:75 ratios. Thus, the somatic mutations across these three tumor mixture ratios have AFs ranging from 5% to 35% for Stage 3 dataset, and 3.75% to 21% for Stage 4 dataset.

### Platinum synthetic tumor data

We downloaded 200*×* Platinum genomes NA12878 and NA12877 and their truth germline variants [17] to construct a virtual tumor and normal pair (ENA accession number PRJEB3246). For the normal, we downsampled NA12877 to 50*×*. For tumor, we constructed three 50*×* in silico mixture samples with 70%, 50%, and 25% tumor purities, by independently downsampling NA12877 respectively at 15*×*, 25*×*, and 37.5*×*, and mixing each with downsampled NA12878 at 35*×*, 25*×*, and 12.5*×*. We use the heterozygous and homozygous variants in NA12878 which are reference calls in NA12877 and are at least 5 bases apart from each other as the truth set for the training and evaluation steps (1,103,285 SNVs and 174,754 INDELs). Thus, depending on the zygosity of the germline variants in NA12878, somatic mutations across these three tumor mixture ratios have AFs ranging from 12.5% to 70%.

We also generated another 50*×* virtual tumor sample by randomly spiking reads from a downsampled (to 50*×* coverage) NA12878 into a downsampled (to 50*×* coverage) NA12877 data at heterozygous and homozygous variant locations in NA12878 which are reference calls in NA12877. The frequencies of spiked reads are sampled from a binomial distribution with means [0.05, 0.1, 0.2, 0.3], thus depending on the zygosity of the variant, the mean somatic mutations AFs ranges from 2.5% to 30%. To avoid ambiguity in the truth set, we only used variants for which the relevant paired-end reads did not overlap any other variants (316,050 SNVs and 46,978 INDELs). This generated a contaminated tumor with reads from NA12878. We also used another independent downsampled (to 50*×*) data for NA12877 as the pure normal.

For both experiments, FastQ files and truth germline variants were downloaded and aligned with BWA-MEM (v0.7.15) [25] followed by Picard MarkDuplicates (v2.10.10) (https://broadinstitute.github.io/picard), and GATK IndelRealigner and Base Quality Score Recalibration (v3.7) [26].

### Real tumor–normal pair data

We used the COLO-829 immortal metastatic malignant melanoma cell line dataset [21, 22] (accession: EGAS00000000052) to assess our approach on real tumor-normal pair data with published lists of validated somatic mutations.

The COLO-829 dataset consists of 80*×* whole-genome sequencing tumor sample and its matched normal blood COLO-829BL sample at 60*×*, with 454 validated somatic SNVs.

The TCGA-AZ-6601 [23, 24] dataset is a whole exome sequencing of a colon adenocarcinoma tumor sample and its matched normal tissue from TCGA. The tumor and normal samples were sequenced at depths of 145*×* and 165*×* respectively. We used 952 validated SNVs in TCGA [27] and COSMIC [28] databases as the ground truth somatic mutations for this sample.

For real data, we compute pseudo-precision as the percentage of predicted somatic mutations that have been called by at least two stand-alone methods, or have been reported as verified somatic mutations in at least two samples of the same cancer type in COSMIC database. We then compute pseudo F1-score based on the harmonic mean of this pseudo-precision and recall.

### Whole-exome and targeted panel data

To assess NeuSomatic on different target enrichment experiments we used whole-exome datasets from the Ashkenazi Jewish trio [19]. We downloaded deep-sequenced (200*×* coverage) whole-exome alignment files for HG003 and HG004 (ftp://ftp-trace.ncbi.nlm.nih.gov/giab/ftp/), along with the high-confidence germline variants. We then used mixtures of random 70*×*, 50*×*, and 25*×* downsamples of HG004 and 30*×*, 50*×*, and 75*×* downsamples of HG003, to construct 70%, 50%, and 25% pure tumor samples, respectively. We also constructed a 95% pure normal by mixing 95*×* HG003 and 5*×* HG004 downsampled alignments. For our analysis, we used Agilent SureSelect Human All Exon V5 BED file. The ground truth somatic mutations were identified similar to the Platinum synthetic tumor data (11,720 SNVs, 878 INDELs). Depending on the zygosity of the germline variants in HG004, somatic mutations across these three tumor mixture ratios have AFs ranging from 12.5% to 70%

For validating the performance on target panel, we restricted the above alignment and truth data to Illumina’s TruSight inherited disease panel bed file (216 SNVs, 5 INDELs). We only evaluated the performance on SNVs due to the limited number of true INDELs in the target panel region.

### PacBio data

For long-reads analysis, we downloaded the high-confidence germline variants for HG002 sample (ftp://ftptrace.ncbi.nlm.nih.gov/giab/ftp/) [19]. We built the long-reads error profile using the CHM1 dataset [29] (SRA accession SRX533609). We then simulated a 100*×* pure normal sample using the VarSim simulation framework [30] in combination with the LongISLND in silico long-reads sequencer [31]. Using a set of random somatic mutations, we also simulated a 100*×* pure tumor sample with the same error profile. We used NGMLR (v0.2.6) [32] to align the sequences. We then mixed a 47.5*×* downsample of pure normal alignment and 2.5*×* downsample of the pure tumor alignment to form the 50*×* normal pair with 95% purity, and mixed 40*×*, 35*×*, and 25*×* independent downsamples of normal respectively with 10*×*, 15*×*, and 25*×* downsamples of pure tumor, to construct 50*×* tumor mixtures of 20%, 30%, and 50% purity. We restrict the training set to a 120 megabase region in chromosome 1 (with 39,818 truth somatic SNVs and 38,804 truth somatic INDELs), and the testing set to whole chromosome 22 (with 12,201 truth somatic SNVs and 12,185 truth somatic INDELs). Somatic mutations across the three tumor mixture ratios have AFs ranging from 20% to 50%.

### Candidate mutation preparation

As the first step, we scan tumor read alignments to find candidate locations with evidence of mutations. Many of these positions have either germline variants or erroneous calls made due to the complexity of the genomic region, or sequencing artifacts. We apply a set of liberal filters on the set of candidate locations to make sure the number of such locations is reasonable. In general, for SNVs, we required AF*≥* 0.03 or more than 2 reads supporting the SNV and Phred scaled base quality score larger than 19 (larger than 14 for real WES dataset) as the minimum requirements. For 1-base INDELs, we required AF*≥* 0.02 or more than one read support. For INDELs larger than 1-base, we require AF*≥* 0.03. For the ensemble approach, we also included any somatic mutation detected by other somatic mutation detection methods as input candidate. For the PacBio dataset, we used AF*≥* 0.1 for SNVs and INDELs larger than 1-base, and AF*≥* 0.15 for 1-base INDELs.

For the DREAM Challenge dataset, we excluded variants existing in dbSNP [33] from the input candidates.

### Input mutation matrix

For each candidate position, we prepare a 3-dimensional matrix *M* with *k* channels of size 5 *×* 32 (Figure 1a; Supplementary Figures 1 and 2). The 5 rows in each channel corresponds to 4 DNA bases A, C, G, and T, and the gap character (‘-’). Each of the 32 columns of the matrix represent one column of the alignment.

For each candidate location, we extract the tumor and normal read alignments. As shown in Figure 1a, we then consider the read alignments of tumor and normal sample to the reference as an MSA. To this end, we augment the reference sequence by adding gaps to the reference sequence, when there is insertion in reads. It must be noted that this process does not need any further realignment of the original read alignments of input BAM files, but only restructuring the alignments into MSA format by assigning additional columns wherever insertions has occurred. If there are multiple distinct insertions in multiple reads after a specific position, we consider them as left-aligned sequences and put them in the same set of columns (See for instance insertions of A and C bases in the 9th column of the toy example in Figure 1a). With this read representation, we find the frequency of A/C/G/T/characters in each column and record separate matrices for tumor and normal samples (channels *C*2 and *C*3 in matrix M). In channel *C*1, we record the reference base (or gap) in each column. Channels *C*_*i*_ (4 *≤ i ≤ k*) record other alignment signals in tumor and normal samples, such as coverage, base quality, mapping quality, strands, clipping information, edit distance, alignment score, and paired-end information. For instance, for the base quality channel, we have a matrix of size 5 *×* 32 for each sample which records the average base quality of reads that have a given base (for a given row) in each column. As another instance, for the edit distance channel, we have a matrix of size 5 *×* 32 for each sample which records the average edit distance of reads that have a given base (for a given row) in each column. One channel of matrix *M* is devoted to specify the column where the candidate is located in. In the current implementation, we use total of 26 channels in the stand-alone NeuSomatic-S approach.

For the ensemble extension of NeuSomatic, we also included additional channels to capture features reported by each of the 6 individual methods used. In this implementation, we used 93 additional channels to represent features extracted from other methods, and alignments reported by SomaticSeq. Thus, the ensemble mode of NeuSomatic had 119 input channels for each candidate matrix.

For each candidate location, we report the alignment information in a window of 7 bases around the candidate. We reserve 32 columns to take into account the augmented alignment with insertions. In rare cases where we have a large insertion, 32 columns may not be enough to represent the alignment. For such cases, we truncate the insertions so that we can record at least three bases in the vicinity of the candidate.

### CNN architecture

The proposed CNN (Figure 1b) consists of 9 convolutional layers structured as follows. The input matrices are fed into the first convolution layer with 64 output channels, 13 kernel size and Relu activation followed by a batch normalization and a max-pooling layer. The output of this layer is then fed to a set of four blocks with shortcut identity connection similar to ResNet structure. These blocks consist of a convolution layer with 3 kernels followed by batch normalization and a convolution layer with 5 *×* 5 kernels. Between these shortcut blocks, we use batch normalization and max-pooling layers. The output of final block is fed to a fully connected layer of size 240. The resulting feature vector is then fed to two softmax classifiers and a regressor. The first classifier is a 4-way classifier that predicts the mutation type from the four classes of non-somatic, SNV, insertion, and deletion. The second classifier predicts the length of the predicted mutation from the four categories of 0, 1, 2, and *≥* 3. Non-somatic calls are annotated as zero size mutations, SNVs and 1-base INDELs are annotated as size 1, while 2-base and *≥* 3 size INDELs are respectively annotated as 2 and *≥* 3 size mutations. The regressor predicts the column of the mutations in the matrix, to assure the prediction is targeted the right position and is optimized using a smooth L1 loss function.

The CNN has less than 900K parameters which enables us to have a highly efficient implementation by using large batch sizes. The whole-genome training process took *∼*8 hours on a machine with 8 Tesla K80 Nvidia GPU’s.

### CNN training

For DREAM Challenge, Platinum, and target enrichment datasets, we randomly split the genomic regions to 50% training and 50% testing sets. For hyper-parameter tuning, we only used 10% of the genome in the DREAM Challenge Stage 3 experiment and used the derived parameters in all other experiments. For the PacBio dataset, we trained NeuSomatic on a 120 megabase region on chromosome 1, and tested it on all of chromosome 22.

The DREAM Challenge dataset has 15,507 somatic mutations for Stage 3 and 30,462 somatic mutations for Stage 4. For better network training, we spiked in *∼*95K more SNVs and *∼*95K more INDELs with similar AF distributions to the original DREAM data into the tumor samples of Stages 3 and 4 using BAMSurgeon [18].

We trained the network using a batch size of 1000 with SGD optimizer with learning rate of 0.01, and momentum of 0.9, and we multiplied the learning rate by 0.1 every 400 epochs.

Since, in general, the input candidate locations have significantly more non-somatic (reference or germline) calls than true somatic mutations, in each epoch, we use all the true somatic mutations in the training set and randomly selected non-somatic candidates with twice the number of the true somatic mutations. We used a weighted softmax classification loss function, to balance for the number of candidates in each category. For DREAM Challenge data, since we added more synthetic mutations in the training set, we boosted the weight for the non-somatic category to achieve higher precision on test set.

For assessing synthetic target enrichment datasets, we used whole-exome and whole-genome data as the training set.

To test on real WGS sample COLO-829, we used models trained on DREAM Challenge Stage 3 with 70% purity for SomaticSeq and NeuSomatic. For the real WES sample TCGA-AZ-6601, we prepared a training set using data from another TCGA WES dataset, TCGA-AZ-4315 [27]. We mixed the tumor and normal alignments from this dataset and split the mixture into two equal alignments. We then used one alignment as the pure normal and spiked in *∼*91K random SNVs and *∼*9K random INDELs into the other alignment using BAMSurgeon to generate a synthetic tumor sample for training. We used models trained on this synthetic tumor-normal WES dataset to test NeuSomatic and SomaticSeq on the real WES dataset, TCGA-AZ-6601.

### Training on multiple purities

For more generalizable training, we combine the generated training input matrices from different tumor purities in the DREAM Stage 3 dataset, and used the combined set for training the network. We then applied the trained model on each of the individual purities.

### Other somatic mutation detection algorithms

We used Strelka2 (v2.8.4), Mutect2 (v4.0.0.0), SomaticSniper (v1.0.5.0), MuSE (v1.0rc), VarDict (v1.5.1), VarScan2 (v2.3.7), and SomaticSeq (v2.7.0) somatic mutation detection algorithms in our analysis, with their default settings.

We used VarDict as an alternative approach to NeuSomatic on PacBio data. To enable detecting somatic mutations on high-error rate long reads, we used VarDict with “`-m 10000 -Q 1 -q 5 -X 1 -c 1 -S 2 –E 3 -g 4 -k 0` “parameter settings. Besides, as in NeuSomatic, we used AF*≥* 0.1 for SNVs and AF*≥* 0.15 for INDELs.

To train SomaticSeq, we also followed the same 50% train/test region splitting as used for NeuSomatic. For the precision-recall analysis, somatic mutations were sorted based on the confidence or quality scores assigned by each tool. For MuSE, we used the tier assignments as the sorting criterion. For VarDict, VarScan2, MuTect2, Strelka2, and SomaticSniper we respectively use SSF, SSC, TLOD, SomaticEVS, and SSC values reported in the VCF file for sorting. For SomaticSeq and NeuSomatic, we use the somatic mutation quality score in the QUAL field. NeuSomatic reports quality scores for the predicted somatic mutations based on the probabity of predictions by CNN.

To analyze performance on real samples, we used the PASS somatic calls from different methods (For VarDict we restrict to calls with StrongSomatic status). For NeuSomatic, we used 0.97 as the quality score threshold for WGS and 0.6 for WES.

### Computational complexity

For whole-genome data, scanning 50*×* tumor and normal alignments to find candidates, extracting features, and preparing the input matrices can take *∼*5.5 hours on a dual-14 core Intel Xeon^®^ CPU E5-2680 v4 2.40GHz machine. The whole-genome training process can take *∼*8 hours on a machine with 8 Tesla K80 Nvidia GPU’s (90 Seconds per epoch of size 580,000). Depending on the cutoff AF on candidate somatic mutations, computing the network predictions for the candidate mutations on a 50*×* whole genome data can take *∼*50 minutes (with AF cutoff of 0.05, 3.5M candidates) to *∼*2.5 hours (with AF cutoff of 0.03, 11M candidates) with 8 Tesla K80 Nvidia GPUs.

## Acknowledgements

We thank Li Tai Fang, Yunfei Guo, Lijing Yao, Betty Ulitsky, Jian Li, Anoop Grewal, Daniel Kogan and Michael Braverman from Roche Sequencing Solutions for fruitful discussions and proofreading the manuscript.

## Author contributions

S.M.E.S., M.M., and H.Y.K.L conceived the study, reviewed the analysis, and wrote the manuscript. H.Y.K.L designed the study. M.M. and H.Y.K.L. supervised the study. S.M.E.S designed and developed the NeuSomatic algorithm and its software. R.L. and S.M.E.S developed the algorithm for scanning the genome, preparing the augmented alignments, and extracting alignment features. S.M.E.S performed the overall analysis. B.L., S.M.E.S., and M.M. performed the long-read analysis. All authors reviewed and edited the manuscript.

## Competing financial interests

S.M.E.S, M.M., and H.Y.K.L. have filed a patent application on this invention.

## Code Availablity

NeuSomatic is written in Python and C++. Its deep learning framework is implemented using PyTorch 0.3.1 to enable GPU application for training/testing. NeuSomatic is open-source and available at https://github.com/bioinform/neusomatic.

